# Genomic differences and species delimitation: a case for two species in the zoonotic cestode *Dipylidium caninum*

**DOI:** 10.1101/2023.02.23.529708

**Authors:** Jeba R J Jesudoss Chelladurai, Aloysius Abraham, Theresa Quintana, Vicki Smith, Deb Ritchie

**Affiliations:** Department of Diagnostic Medicine/Pathobiology, College of Veterinary Medicine, Kansas State University, Manhattan, KS, USA 66506; Department of Biotechnology, Alagappa University, Karaikudi, India 630003

## Abstract

*Dipylidium caninum* (Linnaeus, 1758) is a common zoonotic cestode of dogs and cats worldwide. Previous studies have demonstrated the existence of largely host associated canine and feline genotypes based on infection studies, genetic differences at the nuclear 28S rDNA gene and complete mitochondrial genomes. There have been no comparative studies at a genome-wide scale. Here, we sequenced the genomes of a dog and cat isolate of *Dipylidium caninum* from the United States using the Illumina platform and conducted comparative analyses with the reference draft genome. Complete mitochondrial genomes were used to confirm the genotypes of the isolates. *D. caninum* canine and feline genomes generated in this study had mean coverage depths of 45x and 26x and an average identity of 98% and 89% respectively when compared to the reference genome. SNPs were 20 times higher in the feline isolate. Comparison and species delimitation using universally conserved orthologs and protein coding mitochondrial genes revealed that the canine and feline isolates are different species. Data from this study builds a base for future integrative taxonomy. Further genomic studies from geographically diverse populations are necessary to understand implications for taxonomy, epidemiology, veterinary clinical medicine, and anthelmintic resistance.

## Introduction

*Dipylidium caninum* (Linnaeus, 1758) is a cosmopolitan cestode belonging to the family Dipylidiidae of the order Cyclophyllidea. It is capable of infecting domestic dogs, domestic cats [1], wild carnivores [2] and humans [3]. Taxonomically, it is currently accepted that *D. caninum* occurs as two distinct, host-associated genotypes named the *“D. caninum* canine genotype” and *“D. caninum* feline genotype”.

Infection in definitive hosts is acquired through the ingestion of the cysticercoid stage present within insect intermediate hosts - adult fleas of the genus *Ctenocephalides, Pulex* or adult lice of the genus *Felicola* [4–7]. Cysticercoids are released within the intestines and develop into a scolex followed by development and maturation of immature and later gravid proglottids. Gravid proglottids are released into the intestines and pass out in feces. Gravid proglottids released into feces may move around the perineal region or bedding/furniture. These may occasionally cause pruritis of the peri-anal region resulting in scooting behavior in dogs. Egg packets containing 5 – 30 oncospheres are released by proglottid disintegration or active extrusion. The oncospheres are ingested by the larvae/juveniles of the intermediate hosts. Thus, the life cycle is indirect and *D. caninum* infection in dogs and cats is often associated with an infestation with fleas or lice.

*D. caninum* has a moderately broad host-specificity. The presence of two distinct, host-associated genotypes was first demonstrated in phylogenetic analyses of the partial 28S genes [6], partial mitochondrial 12S genes [7], and complete mitochondrial genomes [8]. In a majority of naturally infected cases, dog and cat hosts are infected with their respective genotypes. Cat lice and flea-derived *D. caninum* from Malaysia belonged to the feline genotype based on 12S analyses[7]. Seven of 9 cat derived *D. caninum* isolates from the United States belonged the feline genotype at the 28S gene (genotypes from 2 isolates could not be determined) [8]. Dog flea-derived *D. caninum* from across Europe belonged to the dog genotype (100%), whereas 95.1% of cat flea-derived D. caninum from the same area belonged to the feline genotype based on 28S analyses [8]. Praziquantel resistant *D. caninum* samples from the United States that were obtained from dog feces were found in the canine genotype clade at both the partial 12S and 28S genes, showing host-association [9].

The distinction was further substantiated by *in vivo* experimental studies. Prepatent periods of the infection were shorter and lifespans longer when host associated genotypes infected the appropriate host [10]. There was no evidence of *in vivo* hybridization between the feline and canine genotypes. Host specificity and preference breaks down in only in 2-10% of natural infections [8]. In these cases, the feline genotype can be recorded in dogs or in fleas isolated from dogs and vice versa. Experimentally, dogs are “permissive” to infection by the feline genotype and cats are “permissive” to infection by the canine genotype [10]. Despite the permissivity, genotypes show biological adaptation with improved longevity and shorter prepatent periods in their respective hosts. Wildlife – hyaenas, red foxes - appear permissive to infections with the feline genotype [2,11]. Humans are also permissive to infection by the feline genotype [12].

No comparisons of canine and feline genotypes have been made at the whole genome level yet. If the two genotypes are two species, clinical implications exist for veterinarians who treat and control infections in the face of increase in anecdotal reports of praziquantel resistance. Genetic differences at specific nuclear and/or mitochondrial genes and at the genome level are useful for differentiating species through species delimitation algorithms. Species delimitation has been used to resolve taxonomic conundrums in cestodes like *Mesocestoides* [13] and in other eukaryotes [14]. Recently, universal single copy orthologs (USCOs) have been demonstrated to provide high resolution to differentiate between closely related species [15]. USCO genes from cat isolates of *D. caninum* have not been described yet. USCO genes from a dog isolate of *D. caninum* from China are available along with a draft nuclear genome [16], which can serve as a reference in comparative studies.

Our objectives in this study were to sequence the genomes of *D. caninum* isolated from a dog and a cat from the United States using the Illumina platform and to compare them to the reference *D. caninum* genome isolated from a dog in China [16]. We hypothesized that the genomes of *D. caninum* canine isolates would be similar despite the geographical distance of the sites of isolation, and that *D. caninum* feline isolates would have significant differences. Mitochondrial genomes were used to confirm the identity of dog and cat isolates. Genomes were compared and a set of single copy orthologues were used in phylogenetic and species delimitation analyses.

## 2. Materials and methods

### 2.1 Parasite material and sequencing

Feces of a dog in Florida, USA naturally infected with *Dipylidium caninum* was collected (isolate: Canine FL1). *Dipylidium caninum* proglottids were isolated by mixing the feces with water and sieving through a 1 mm sieve. Proglottids were removed from the sieve using forceps, thoroughly washed with 1x phosphate-buffered saline and identified using egg packet and proglottid morphology. *D. caninum* proglottids passed by a cat in Kansas was isolated from the perineal area, washed thoroughly in phosphate-buffered saline and identified by morphology of egg packets and proglottids (isolate: Feline KS1).. All proglottids were stored in 70% ethanol at −20°C until DNA extraction.

Genomic DNA was extracted using the DNeasy Blood and Tissue Kit (Qiagen, Valencia, CA) using the manufacturer’s protocol. An RNase treatment was performed to remove co-purified RNA. Sample quantity was assessed using a Qubit fluorometer (Thermo Fisher Scientific, Waltham, MA). Sample quality was assessed using an Agilent 5400 Bioanalyzer (Agilent Technologies Inc., California, USA). Genomic DNA was first fragmented using Covaris in the 350 bp mode. Library preparation was performed with NEB Next Ultra DNA Library Prep Kit (Ipswich, MA) following manufacturer’s instructions. Briefly, blunt ends fragments were generated by end-repairing 3’or 5’overhangs of double-stranded DNA (dsDNA) fragments followed by 3’ dA-tailing, index adapter ligation and size selection using SPRIselect beads. This was followed by PCR enrichment of the adaptor ligated library. Samples were pooled and sequenced on Illumina HiSeq 4000 sequencer for 150 bp read length in paired-end mode, with an output of 17.9 million paired end reads for the cat sample and 18.9 million paired end reads for the dog sample. Raw data is publicly available on NCBI Bioproject Accession: PRJNA768484; Sequence Read Archive (SRA) Accessions: SRX12485835, SRX12485836.

### 2.2 Assembly, mapping and variant analysis

FastQC (version 0.11.9) [17] was used to assess sequence quality before and after adapter trimming with Trimmomatic (version 0.36) [18]. Mitochondrial genomes were assembled using Novoplasty (Version 4.3.1) [19], annotated with MITOS2 [20] and manual curation. Mitochondrial genomes were submitted to GenBank (Accession numbers: OK523384.1, OK523385.1). Identity of the genotypes were confirmed by BLAST [21] comparisons to previously described mitochondrial genomes [8,22,23].

Raw reads from the *D. caninum* Canine FL1 and Feline KS1 isolates were mapped to the previously described draft reference genome (Assembly Accession number: GCA_017562135.1) using BWA-MEM2 (Version 2.2.1) [24]. Coverage of the genomes was assessed using Qualimap (Version 2.2.2) [25] and bamCoverage (Version 3.5.1) [26], and then visualized with IGV-Web [27]. Variant analysis was conducted with DeepVariant [28]. Reference guided assembly of draft genomes of the two isolates from this study was created with bcftools [29]. De novo assemblies were created with SPAdes (Version 3.15.4) [30]. Similarities and one-to-one comparisons between the genomes were conducted using dnadiff [31]. Assembly, mapping and variant analysis was conducted on Galaxy servers [32].

The number of variants at each scaffold of the reference genome was plotted using vcfR (Version 1.13.0) [33] and *ggplot2* [34] in R. Summary statistics of the variant analysis was plotted with *ggplot2* [34] in R.

### 2.3 Benchmarking Universal Single Copy Orthologs

Genome completeness was assessed with BUSCO (Version 5.2.2) with metazoan lineage parameters in genome mode with the metaeuk predictor [35]. A total of 954 BUSCO groups were searched for each draft genome. Complete BUSCO genes were extracted from the draft assemblies from this study and the reference genome of *D. caninum* using bedtools (Version 2.30.0) [36] and parsed with biopython (Version 1.80) [37]. BUSCO completeness and genes present in the BUSCO sets between the three genomes were visualized with *ggplot2* and *ggvenn* in R (Version 4.1). For each gene, sequences were aligned with MAFFT (Version 7.487) [38] and trimmed with trimal (Version 1.2) [39]. Pairwise genetic distance matrices of the 503 complete BUSCO genes present in the three genomes were calculated using the TN93 model [40] with *apex* (Version 1.0.4) [41]. A heatmap of the calculated distances were created in *ComplexHeatmap* (Version 3.16) [42]. Principal component analysis of the SNPs present in the 503 BUSCO genes were analyzed with *adegenet* (Version 2.1.1) [43] and plotted with *ggplot2* [34].

### 2.4 Phylogenetic and species delimitation analyses

The 3 sets of BUSCO genes obtained above and BUSCOs from 14 other cestode assemblies from GenBank were parsed with biopython (Version 1.80). Only BUSCO genes (128 genes) present in all 17 assemblies were used in the phylogenetic analysis. For each of 128 genes, sequences were aligned with MAFFT (Version 7.487) [38] and trimmed with trimal (Version 1.2) [39]. A concatenated supermatrix and gene partition file of the 128 BUSCO genes were created using *phylotools* (Version 0.2.2) [44] in R. Maximum likelihood phylogenetic reconstruction of the concatenated supermatrix was performed with IQtree2 (Version 2.1.0) [45] with ultrafast bootstrap approximation [46], using ModelFinder [47] to determine the best-fit model for each gene in the supermatrix (that is, partition model) [48], according to Akaike information criterion (AIC) scores and weights. The Diphyllobothridean clade represented by *Schistocephalus solidus* and *Spirometra erinaceieuropaei* was used as the outgroup. Species delimitation analyses of the trees were carried out using Bayesian PTP [49] and ASAP with the 2-parameter Kimura-80 model [50].

GenBank records of complete mitochondrial genomes of cestodes of veterinary interest were obtained. Nucleotide sequences of mitochondrial protein coding genes (12 genes) were parsed with the GenBank Feature Extractor [36]. A concatenated supermatrix and partition file of the 12 protein coding genes was created in *phylotools* (Version 0.2.2) [44] in R. Maximum likelihood phylogenetic reconstruction of the concatenated supermatrix was performed with gene partitions as described for BUSCO genes. The mitochondrial genome of *Schistosoma mansoni* was used as the outgroup. Species delimitation analyses of the mitochondrial genome dataset were carried out as described for BUSCO genes.

## 3. Results

### 3.1 Identity confirmed with complete mitochondrial genomes

Proglottids were identified as *Dipylidium caninum* by morphology. To confirm host-associated genotype identity, complete mitochondrial genomes generated from the Illumina dataset were used. The mitochondrial genomes from the *D. caninum* Canine FL1 and Feline KS1 genomes generated in this study were 14,296 bp and 13,598 bp long respectively. The difference in length is in agreement with previously described mitochondrial genomes lengths [8,22,23]. The complete mitochondrial genome of the *Dipylidium caninum* Canine FL1 isolate (Accession number: OK523384.1) had 97.65% - 99.82% identity with the genomes of canine isolates described earlier [22,23]; however, Identity with the mitochondrial genomes of the feline isolates (This study, [8] were only 84.25% to 86.21% (Table S1). The mitochondrial genome of the *Dipylidium caninum* Feline KS1 isolate from this study had 99.51% identity with the mitochondrial genome of the feline isolate previously described [8]. Thus, the two isolates sequenced in this study were identified to belong to the host-associated genotypes of their host of origin.

### 3.2 Quality summary of the datasets

*D. caninum* Canine FL1 and Feline KS1 isolates from this study generated 18,886,666 and 17,928,712 raw reads (SRA Accessions: SRX12485835, SRX12485836), of which 18,827,832 (99.68%) and 17,870,800 (99.71%) reads were paired, with index trimmed mean insert lengths of 137.4 bp and 138.6 bp respectively. Trimmed reads were aligned with the *D. caninum* canine reference draft genome (Assembly Accession number: GCA_017562135.1) [16], which has 1686 scaffolds (Figure 1). Average depth of coverage of the *D. caninum* canine FL1 and feline KS1 genomes generated in this study were 46.5x and 25.8x with a GC% of 47.71% and 47.33% respectively. Draft genomes generated using the reference guided assembly generated assemblies that were 108.97 Mb and 108.99 MB for the canine and feline isolates respectively; de novo assemblies were 120.43 Mb and 133.53 Mb respectively.

**Figure 1.**
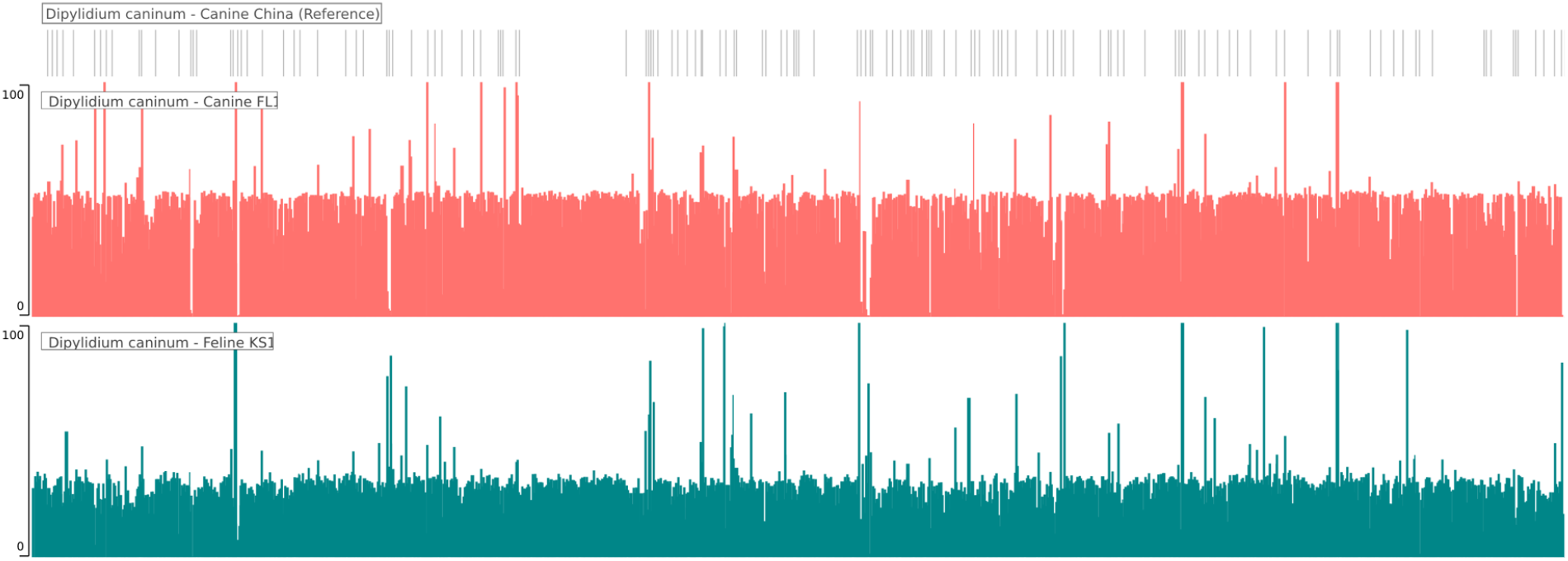
Whole genome coverage plot using *Dipylididum caninum* canine isolate FL1 (red-orange) and *Dipylididum caninum* feline isolate KS1 (teal) mapped to the reference *Dipylididum caninum* draft genome (grey). Average depth of coverage was 46.5X for the canine FL1 isolate and 25.8x for the feline KS1 isolate.

### 3.3 Genomic differences and variation

*D. caninum* Canine FL1 and Feline KS1 isolates from this study were compared to the scaffolds of the reference genome and genetic variants determined from mapped reads. Total number of variants that passed quality checks were 204,341 and 3,495,868 in the *D. caninum* canine FL1 and feline KS1 isolates respectively. These variants were found across 741 and 905 scaffolds of the reference genome in the comparisons. Mapping of variant counts across the scaffolds of the reference genome is shown in Figure 2A. Mean number of variants per scaffold were 275.8 (median 21) and 3863 (median 110) in the *D. caninum* Canine FL1 and Feline KS1 isolates respectively (Figure 2B). To account for the length of each reference scaffold, mean number of variants per 1000 base pairs of reference scaffold was 1.31 (median 0.88) and 10.98 (median 4.54) in the *D. caninum* Canine FL1 and Feline KS1 isolates respectively (Figure 2C). There were 3.3 million biallelic SNPs in the Feline KS1 isolate when compared to the reference genome, which was higher than the 164,000 SNPs in the Canine FL1 isolate (Figure 3A). Biallelic insertions and deletions were also higher in the Feline KS1 isolate than in the Canine FL1 isolate. Transition/transversion ratio was 1.95 and 1.96 in the Canine FL1 and Feline KS1 isolates respectively (Figure 3B). Biallelic base changes from reference are shown (Figure 3C).

**Figure 2.**
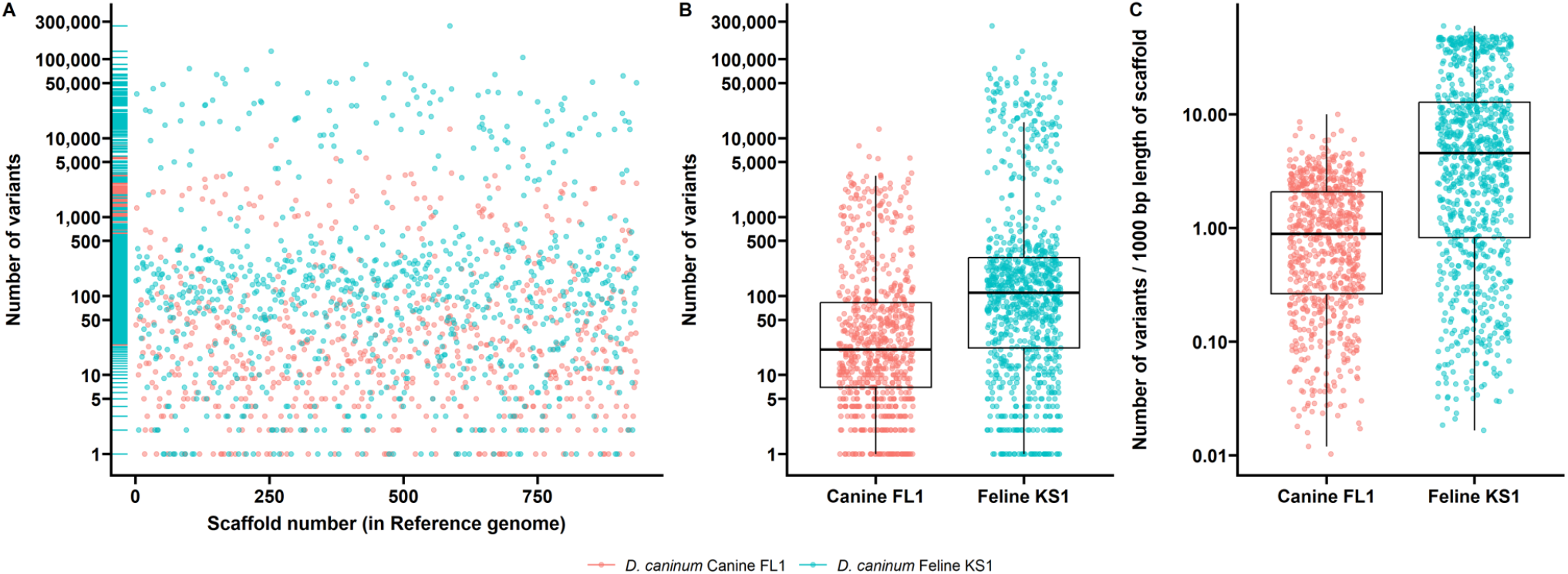
Genetic variants in the genomes D. *caninum* canine FL1 and D. *caninum* feline KS1 compared to the reference genome of *D. caninum*. (A) Scatterplot showing the number of genetic variants (SNPs, Indels) at each scaffold location. (B) Boxplots showing the distribution of the number of variants in each genome.

**Figure 3.**
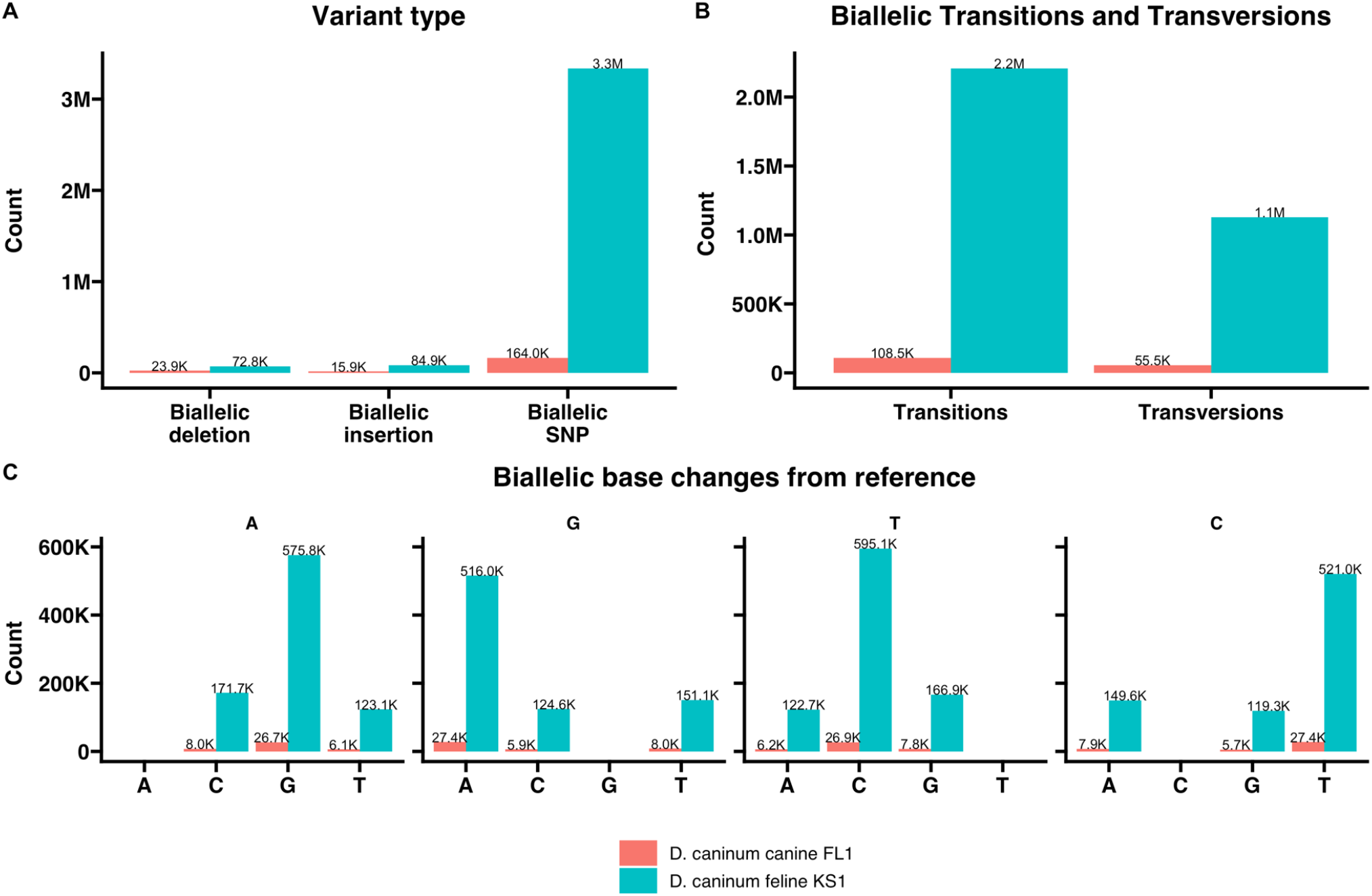
Summary of genetic variants in the comparative analyses.

Draft genomes of the *D. caninum* Canine FL1 and Feline KS1 isolates from this study were compared with each other and the reference genome. The Canine FL1 isolate had an average identity of 99.01% when compared to the reference genome in one-to-one alignments, whereas the Feline KS1 isolate had an average identity of 88.89% when compared to the reference genome. The draft genomes of the *D. caninum* Canine FL1 and Feline KS1 isolates from this study were 88.98% identical. Thus, a ~11% difference exist between the genomes of D. caninum canine and feline genotypes.

### 3.4 BUSCO statistics and comparisons

Complete and single copy BUSCO genes from the Canine FL1 and Feline KS1 isolates were 608 (63.7%) and 552 (57.9%) in number respectively out of the 954 tested in the metazoan lineage. BUSCO genes in the reference assembly (Assembly Accession number: GCA_017562135.1) was 602 (63.1%) (Figure 4). There were 503 orthologs present in all three assemblies. Nucleotide sequences of BUSCO genes of the two isolates from this study were compared with each other and the reference genome. BUSCOs from the Canine FL1 isolate had an average identity of 99.43% when compared to BUSCOs from the reference genome in one-to-one alignments, whereas BUSCOs from the Feline KS1 isolate had an average identity of 91.36% when compared to the BUSCOs from the reference genome. BUSCO genes from the Canine FL1 and Feline KS1 isolates from this study were 91.53% identical.

**Figure 4.**
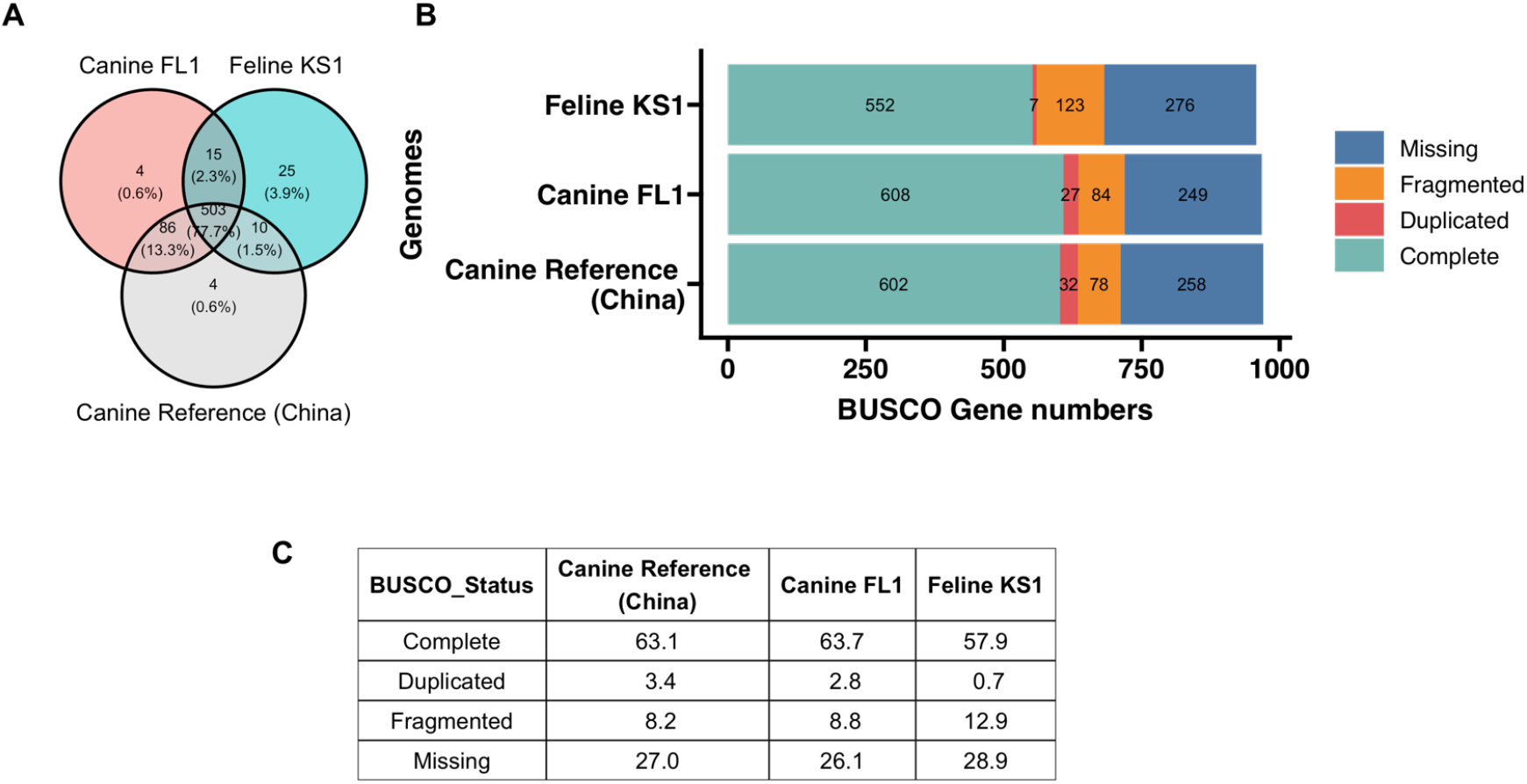
(A) Venn diagram of overlapping complete BUSCO genes among the three genomes. (B) BUSCO assessment results of the three genomes of *D. caninum*. (C) Percentages of BUSCO genes.

Pairwise genetic distances were calculated at the BUSCO genes (503 genes) that were present in all three assemblies and mapped using a heatmap. Genetic distances between genes of the two compared canine isolates were closer than the distances between genes of the feline isolate and the canine isolates (Figure 5).

**Figure 5.**
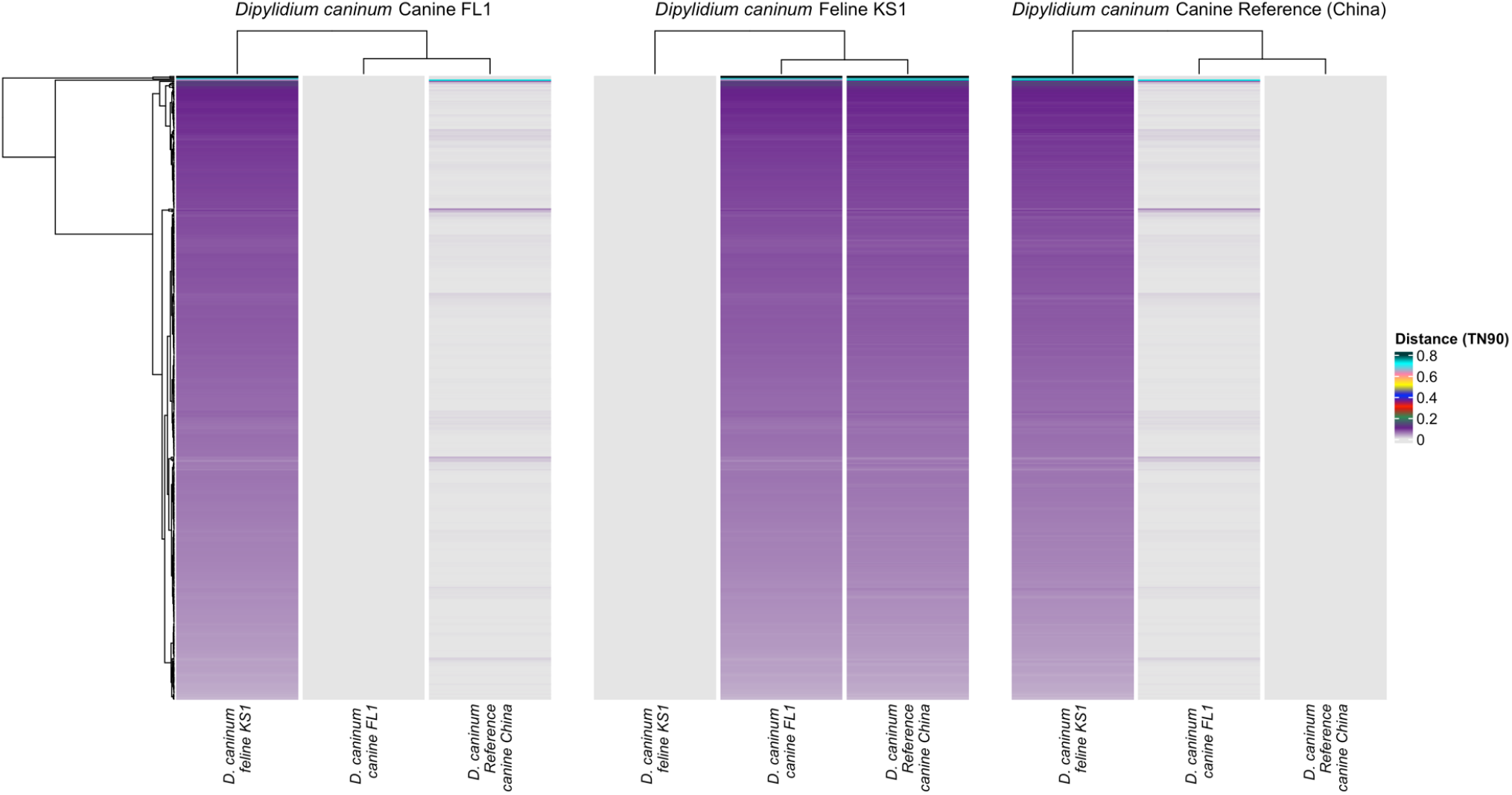
Heatmap of distance matrices at 503 complete BUSCO loci calculated with the Tamura-Nei 1993 model. Each line in the heatmap represents one gene. Color legend indicates genetic distance. (A) Distances of the *D. caninum* canine FL1 BUSCO genes with itself and the other *D. caninum* genomes. (B) Distances of the *D. caninum* feline KS1 BUSCO genes with itself and the *D. caninum* canine genomes. (C) Distances of the *D. caninum* canine reference BUSCO genes with itself and the other *D. caninum* genomes.

Additionally, a principal component analysis was performed to study the relationships between BUSCO genes from the three genomes. SNPs in the 503 shared BUSCO genes were used in the analysis. The first principal component explained 96.82% of the variation and the second principal component explained 3.18% of the variation (Figure 6). The variances between the two canine isolates were similar with overlapping 95% confidence intervals. There was no overlap in the 95% confidence intervals of the feline isolate and both the canine isolates. Thus, *D. caninum* canine and feline genotypes are distinct at BUSCO genes.

**Figure 6.**
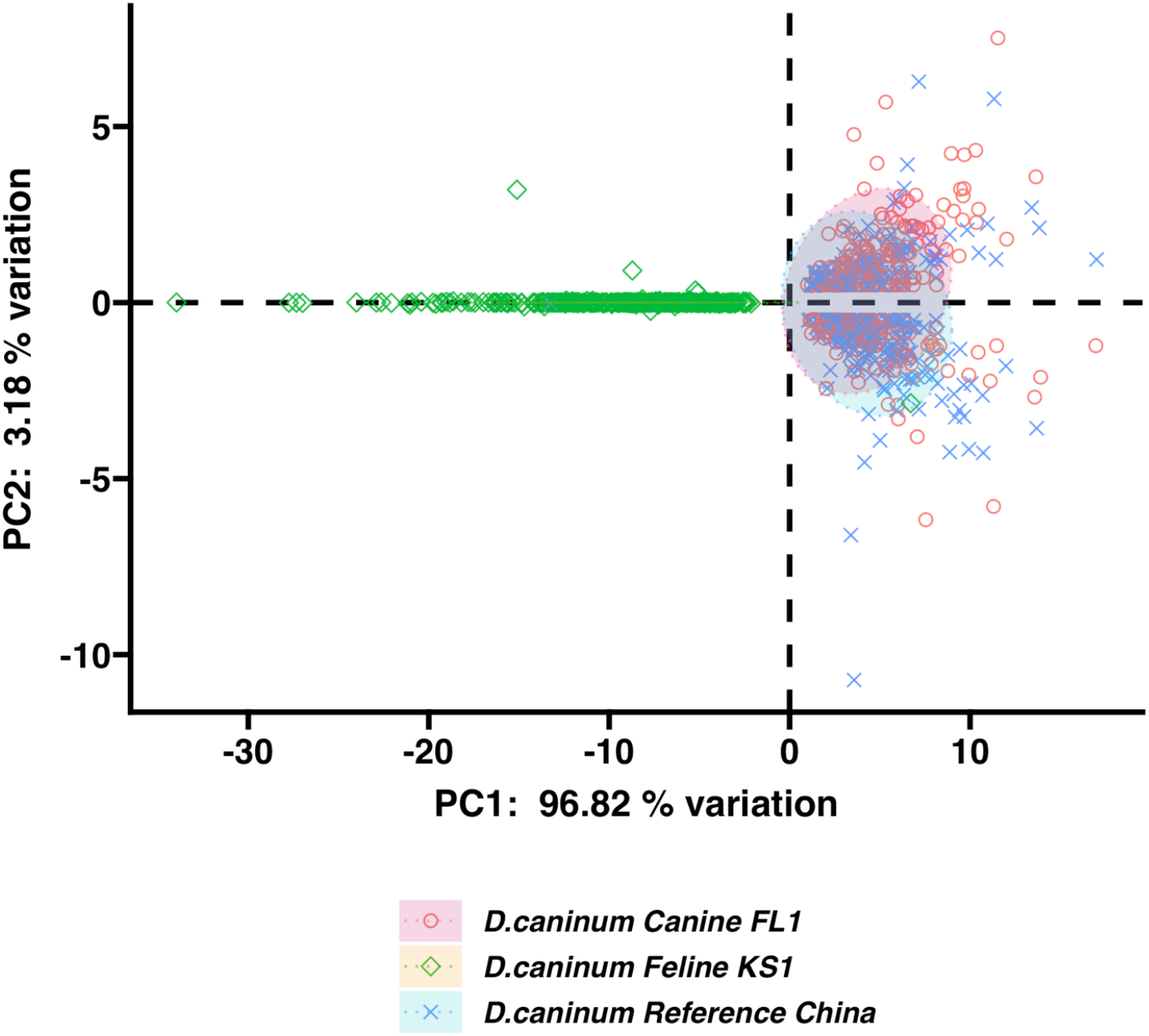
Principal component analysis score plots of SNPs in the 503 BUSCO genes. 95% confidence intervals are shown as ellipses.

### 3.5 Phylogenetic and species delimitation analysis

To understand phylogenetic relationships, a dataset of BUSCO genes from cestode genomes available in GenBank and those from this study was created. Full length, complete genes that were present in all 17 genomes numbered 128 genes. These complete BUSCO genes were concatenated to form a supermatrix of 297,029 nucleotide positions with gene partitions for maximum likelihood phylogenetic analysis, with gene appropriate models for each gene in the supermatrix. The analysis determined 168,880 sites to be parsimony-informative for the maximum likelihood analysis and 75,481 sites to be invariant. The canine isolate of *Dipylidium caninum* in this study formed a monophyletic clade with the canine isolate from China with high statistical support (100%), while the feline isolate KS1 formed a distinct branch (Figure 7). Species delimitation analysis of the BUSCO dataset and tree were carried out. Both the PTP Bayesian solution and ASAP ascending hierarchical clustering solution (Figure 8 colored branches, Figures S2 and S3) identified *D. caninum* Feline KS1 as a species, distinct from the monophyletic clade of the two *D. caninum* canine isolates. The clade with the *D. caninum* canine isolates is supported as a distinct species.

**Figure 7.**
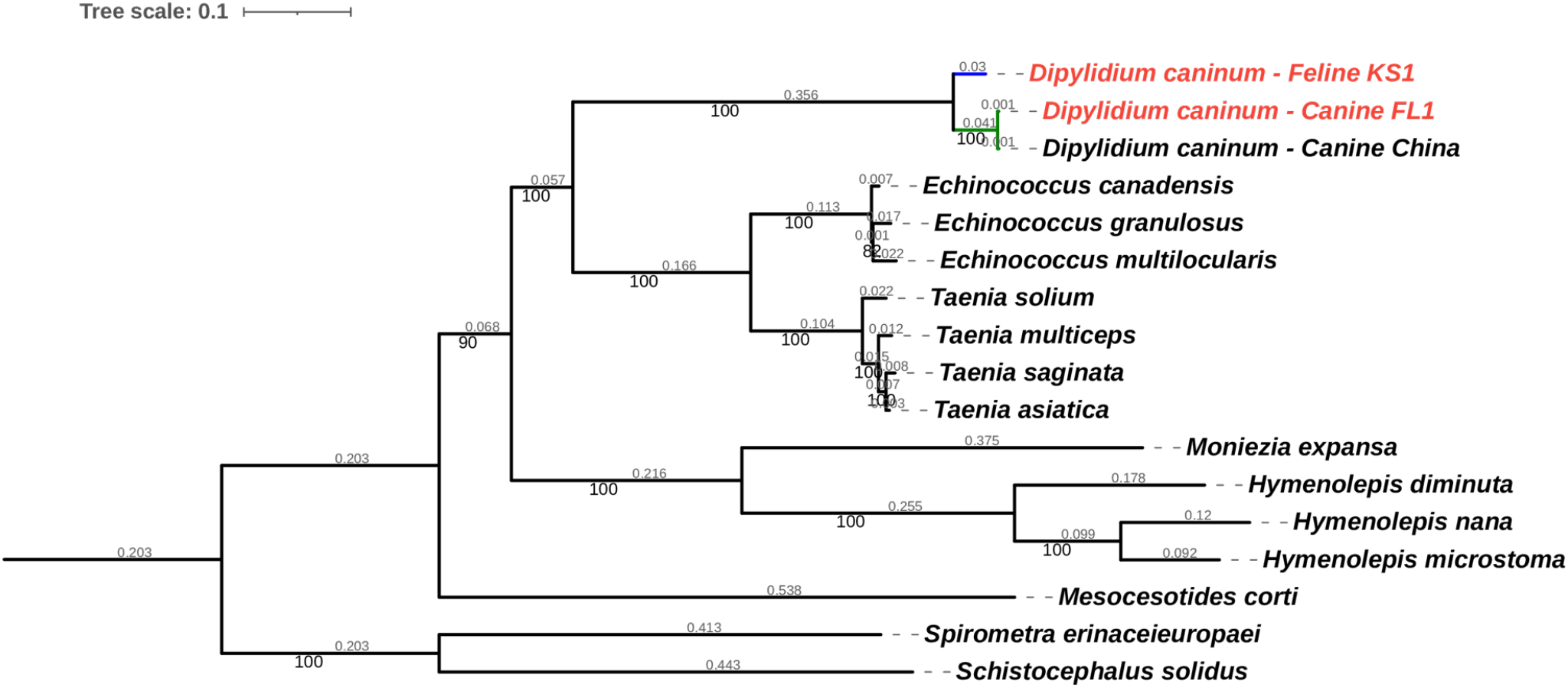
Maximum likelihood phylogenetic trees of 128 BUSCO genes of *Dipylidium caninum* genomes from this study and cestode genomes derived from GenBank, constructed using IQ Tree with gene specific partition models. Accession numbers for genomes derived from GenBank and used in the BUSCO analysis are available in Table S2. Genomes from this study are highlighted in red. Species delimitation of the genus *Dipylidium* is highlighted.

**Figure 8.**
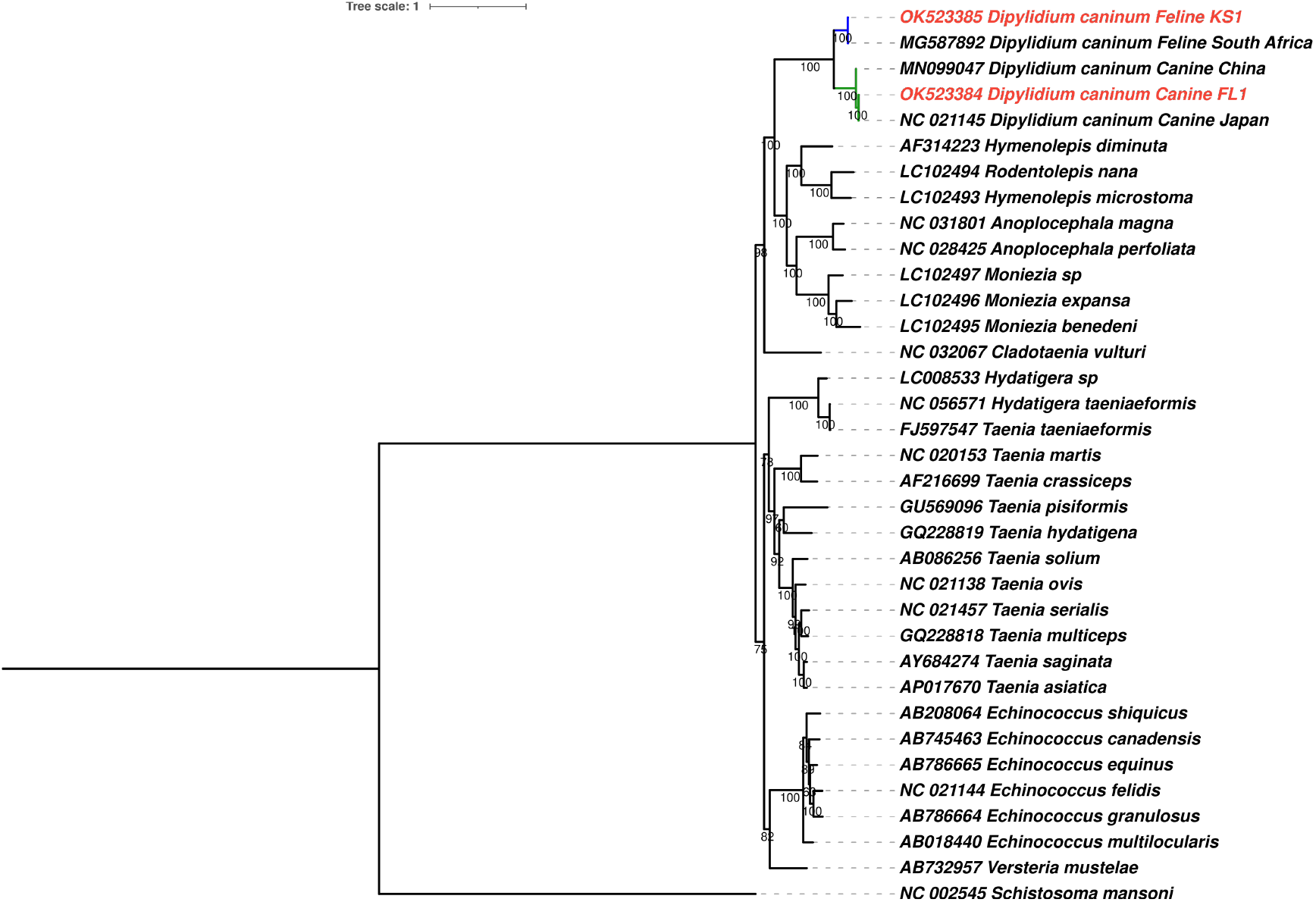
Maximum likelihood nucleotide phylogenetic tree of 12 mitochondrial protein coding genes of *Dipylidium caninum* mitochondrial genomes from this study and those derived from GenBank, constructed using IQ Tree with gene specific partition models. Each leaf of the tree has the GenBank accession and cestode species name. Mitochondrial genomes from this study are highlighted in red. Species delimitation of the genus *Dipylidium* is highlighted. A tree with branch lengths is available in Figure S8.

Additionally, a maximum likelihood phylogenetic tree was constructed from a nucleotide mitochondrial genome dataset, created with the 12 protein coding genes present in the mitochondrial genomes from this study and other cestode mitochondrial genomes available in GenBank (Figure 8). Protein coding genes were concatenated to form a supermatrix of 9948 nucleotide positions with gene partitions for maximum likelihood phylogenetic analysis, with gene appropriate models for each gene in the supermatrix. The analysis determined 5,959 sites to be parsimony-informative for the maximum likelihood analysis and 2,817 sites to be invariant. While the genus *Dipylidium* is monophyletic, there are two distinct clades within it, formed by feline and canine isolates of *D. caninum*. Mitochondrial genomes of *D. caninum* from this study were located within their host-associated clades. Both clades have high statistical support (100%). Bayesian solution-based species delimitation provided support for the two clades to be considered distinct species (Figure 7 colored branches, Figures S5). Hierarchical clustering-based species delimitation provided two solutions when the prior of known species distinctions were applied (Figure S6). The first solution provides evidence for distinct species designations to the host-associated clades, with the feline and canine isolates from this study and prior studies belonging to two distinct species. The second solution suggests that the mitochondrial genome (Accession number: MN099047.1) described from China [22] is a distinct species within the *D. caninum* canine clade. MN099047 could represent a cryptic species. However, species delimitation of the feline isolates was present even in when the second solution is accepted.

## 4. Discussion

*Dipylidium caninum* is a zoonotic cestode that belongs to the family Dipylidiidae within the order Cyclophyllidea. In this study we show for the first time that at the nuclear genome, canine and feline genotypes of *D. caninum* are distinct. In whole genome comparisons, a 11% difference was found to exist between the two genotypes. In representative universal single copy ortholog gene comparisons (503 genes), 8.47-8.64% differences were calculated between the two genotypes. In complete mitochondrial genome comparisons, 13.79-15.83% differences were calculated between the two genotypes. Applying species delimitation criteria to the nuclear and mitochondrial genome data generated in this study and *D. caninum* data from GenBank suggest that the canine and feline genotypes represent two species. Further studies must be undertaken to determine if the two putative species are cryptic or if a taxonomic revision is warranted. Data from this study is available in GenBank (Illumina data - Bioproject Accession: PRJNA768484; mitochondrial genomes - accession numbers: OK523384.1, OK523385.1). We discuss these findings in the context of the historical taxonomy of the species, genomic difference, host specificity, and clinical applications.

*Dipylidium caninum* is the type species within the genus *Dipylidium*, first described in 1863 by Leuckart. Morphological variations in proglottid shape and size, rostellar hook numbers and strobilar length were originally the criteria for describing new species within the genus [51]. Morphology of *Dipylidium* proglottids isolated from dogs and cats have been examined by several authors including Millzner (1926) who compared the morphologies of 1230 cestodes from 28 dogs and 30 cats and described the occurrence of 7 species – *D. caninum* (in a dog), *D. sexcoronatum* (in dogs and cats) *D. gracile* (dogs and cats), *D. crassum* (dogs only), *D. compactum* (cats only), *D. longulum* (cats only) and *D. diffusum* (cats only) [52]. Other host specific *Dipylidium* species described were *D. buencaminoi* (dogs) [53], *D. halli* (cats) *[53], D. carracidoi* (cat) [54], *D. catus* [55] and *D. otocyanois* (bat-eared fox) [56]. However, Venard (1937) used 22 morphological characters to synonymize species described prior to 1937 into three valid species – *D. caninum, D. buencaminoi* and *D. otocyonis*, and suggested that a wide limit of morphological variations be considered normal within the species [57]. Of these three species, scientific literature on the latter two are sparse and *D. caninum* predominates in literature. Several other species previously described in the genus *Dipylidium* have been reassigned to the genera *Diplopylidium* or *Joyeuxiella* within the family Dipylidiiae [58]. Therefore, it is common practice to morphologically identify armed medium-sized cestodes isolated from dogs and cats having a retractable rostellum, double-pored proglottids and eggs present within egg capsules as *Dipylidium caninum*.

Genetic work in the last decade has led to further development in the taxonomy of *D. caninum*. Genetic differences between canine and feline isolates were first observed as sequence variants during the analysis of partial 28S rDNA genes of *D. caninum* cysticercoids [6,8] and based on 28S rDNA, canine and feline host-associated isolates were referred to as canine or feline “genotypes”. Labuschagne et al. used 28S rDNA variation and the complete mitochondrial genomes of canine and feline derived *D. caninum* to propose that the two host associated genotypes be considered distinct species. A 110Mb draft nuclear genome of a canine isolate of *D. caninum* from China is available [16].

The present study is the first to provide additional proof to the two species proposal using whole genome data. Average depth of coverage of the Canine FL1 and Feline KS1 isolates from this study was 46.5x and 25.8x (Figure 1), which was lower than the coverage obtained by [16](200x) due to the different Illumina technologies used. The genomes from this study were mapped against the reference genome to determine the presence and positions of genetic variants such as SNPs, insertions and deletions. Variants were found across the genome in both the Canine FL1 and Feline KS1 isolates from this study (Figure 2A). The total number of variants and the number of variants per 1000 bp of the scaffold were higher in the Feline KS1 genome than in the Canine FL1 genome (Figure 2B, 2C). The number of SNPs in the *D. caninum* Feline KS1 genome compared to the reference was 20.3 times higher than the number of SNPs in the *D. caninum* Canine FL1 genome compared to the reference genome; insertions and deletions were 5.3 times and 3.0 times higher in the *D. caninum* feline KS1 genome. Transitions were higher than transversions in both genomes.

Completeness of the draft genomes were measured using BUSCO genes [35]. BUSCO completeness in the draft Canine FL1 genome (63.1% complete single copy; 3.4% duplicated single copy) was similar to the reference genome (63.5% % complete single copy; 1.3% duplicated single copy) [16]. BUSCO completeness of the draft feline KS1 genome was lower (57.9% complete single copy; 0.7% duplicated single copy). When complete single copy genes were compared between the three genomes, 77.7% were present in all three sets. These were used in further analyses. Based on pairwise genetic distances, BUSCO gene sequences in the two compared canine genotypes are more similar that between the feline genotype and the canine genotypes (Figure 5). Component analysis was applied directly to SNPs in aligned sequence matrices of the shared BUSCO genes. The advantages of this approach are that the structure of the underlying data is not altered and that prior assumptions of relationship are not made [59]. Two distinct clusters with no overlap – the feline genotype cluster and an overlapping larger cluster of the two canine genotypes – were found in the first principal component, accounting for 96.82% of the variation (Figure 6; connected positions at each gene is shown in Figure S1). Thus, the feline and canine genotypes were distinct at >500 universal single copy ortholog genes.

Phylogenetic analysis was performed with BUSCO genes shared by the two genomes in this study and a larger set of cestode genomes. The *D. caninum* canine isolate from this study and the canine reference isolate were found in a monophyletic clade, separated from the feline *D. caninum* isolate (Figure 7). Pairwise genetic distances reflected the phylogenetic structure (Figure S2). Two algorithms were used to analyze species delimitations (Figures S3 and S4) and provided unified evidence that the canine and feline genotypes were distinct species. Interestingly, both the Bayesian and Kimura-80 distances between the feline isolate and the canine isolates were larger than the interspecies distances within the genera *Taenia* and *Echinococcus* (Figures S3 and S4). Given the prior that 4 species of *Taenia* and 3 species of *Echinococcus* were present in the analysis, it is evident from species delimitation analyses that the feline and canine genotypes are distinct species.

Mitochondrial genome analyses and comparisons agreed broadly with the nuclear genome analyses and with the study by [8]. Whole mitochondrial genome differences between the canine and feline genotypes were 13.79%-15.83% (Table S1). Pairwise genetic distance between canine and feline isolates using concatenated 12 protein-coding mitochondrial gene datasets (Tamura-Nei distance = 0.183 - 0.209) were similar to the distances within genera with species which have distinct biology and host preferences, such as *Taenia* and *Echinococcus* (Figure S4). In comparison, pairwise distances between *Taenia solium* and *Taenia serialis* was 0.180 (Tamura-Nei distance) and between *Echinococcus granulosus* and *Echinococcus multilocularis* was 0.174. Distinct clades containing each host associated genotype occurs in the phylogenetic analysis of mitochondrial protein coding genes (Figure 8). This was supported by species delimitation algorithms which provide evidence for the designation of distinct species identities to the canine and feline genotypes based on mitochondrial gene coding sequences (Figure S5 and S6). Interestingly, the mitochondrial genome of an isolate from China (Accession number: MN099047) was considered distinct by one species delimitation algorithm (ASAP; Figure S6) and could represent cryptic species within the dog-specific species of *Dipylidium*. Further studies of mitochondrial genomes of *D. caninum* are warranted to resolve the presence/absence of cryptic species and the range of genetic variation within each host associated *Dipylidium* species.

Nuclear-mitochondrial discordance is evident in the position of the monophyletic clade *D. caninum* in relation to other cestode genera (Figures 7 and 8). In the nuclear BUSCO gene phylogeny, family Dipylidiidae (represented by *D. caninum*) was more closely related to the family Taeniidae (represented by *Echinococcus* spp. and *Taenia* spp.) than to the families Anoplocephalidae (represented by *Moniezia expansa*) and Hymenolepididae (represented by *Hymenolepis* spp.) However, in the mitochondrial gene phylogeny, family Dipylidiidae was more closely related to the families Anoplocephalidae (represented by *Moniezia* spp. and *Anoplocephala* spp.) and Hymenolepididae (represented by *Hymenolepis* spp.) than to the family Taeniidae (represented by *Taenia* spp., *Hydatigera* spp., *Echinococcus* spp. and *Versteria* sp.). The position of the family Dipylidiidae was in agreement with the phylogenetic tree by [60], but different from the tree by [8]. This is likely explained by the fact that [8] used amino acid sequences of the protein coding genes while nucleotide sequences of the protein coding genes were used in this study.

The designation of species status to the canine and feline genotypes has clinical implications for veterinarians in small animal practices and in shelter situations. Based on previous work, the likelihood of encountering host associated *Dipylidium* spp. in pet dogs and cats is higher than the likelihood of encountering non-host associated species (2-10%) [6,8]. Dogs and cats in sympatry in multi-pet households and in shelters may share fleas and *Dipylidium* spp cysticercoids within. Risk of *Dipylidium* spp. infection is higher when pets are flea infested. Further epidemiological work is essential to understand the prevalence of *Dipylidium* spp. cysticercoids in flea populations in different parts of the world and the vectorial capacity of different flea species, since the relative abundance of flea species on pets varies across the world [61,62]. While praziquantel and epsiprantel are useful for treating *Dipylidium* spp. infections in both cats and dogs, veterinarians should be aware that praziquantel resistance has only been reported in canine isolates so far [9].

The small sample size used in this study is a potential pitfall. Although we compared the feline and canine genotypes for the first time, further genetic studies with geographically diverse isolates are essential to increase confidence in the genetic variant calls recorded in this study. The relative low depth of coverage of the isolates is another drawback in the face of the increasing availability of sequencing techniques that provide high coverage [63]. While coverages of 20-30x are common in genomic studies [64], low coverage depths of 4-5x are now being used to detect known and novel variations in larger, more complex eukaryotic genomes [65]. Thus, despite the weaknesses, this study is expected to close a knowledge gap about the difference between the host-associated genotypes of *D. caninum* and can provide a base for integrative taxonomy studies in the future. In light of the new knowledge uncovered in this study, a taxonomic revision of the genus *Dipylidium* may be warranted.

In conclusion, we performed comparative analyses on the nuclear and mitochondrial genomes of dog and cat isolates of *Dipylidium caninum*, representing the canine and feline genotypes. Based on variations, genetic distances, phylogeny and species delimitation from this study, in addition to biological differences demonstrated in experimental studies by [10], there is adequate support for the canine and feline genotypes of *D. caninum* to belong to different species. The nomenclature of the two species must be revisited.

## Funding

This work was supported by the Kansas State University Office of Research Development (University Small Research Grants - 2021 Spring awarded to JJC) and start-up funds provided to JJC by the College of Veterinary Medicine Kansas State University.

## Acknowledgements

The authors would like to acknowledge the Galaxy servers for making servers and software available for analysis. usegalaxy.org is supported by NIH and NSF Grants HG006620, 1661497, and 1929694. usegalaxy.eu is supported by the German Federal Ministry of Education and Research grant 031L0101C and de.NBI-epi. usegalaxy.org.au is supported by Bioplatforms Australia and the Australian Research Data Commons.

## CRediT statement

Conceptualization: J.J.C.; Data curation: J.J.C. and A.A.; Formal analysis: J.J.C. and A.A.; Funding acquisition: J.J.C.; Investigation: J.J.C. and A.A.; Methodology: J.J.C. and A.A.; Project administration: J.J.C.; Resources: J.J.C., T.A.Q., V.S. and D.R.; Software: J.J.C. and A.A.; Validation: J.J.C.; Visualization: J.J.C.; Writing – original draft: J.J.C.; Writing - review & editing: J.J.C., T.A.Q., V.S., D.R. and A.A.;

## Notes

### Competing Interest Statement

The authors have declared no competing interest.

