## Supplemental files for "Genomic differences and species delimitation: a case for two species in the zoonotic cestode *Dipylidium caninum*"

Table S1. BLAST results of complete mitochondrial genomes

| Accession<br>Length<br>Host<br>Reference | OK523384.1<br>14296 bp<br>Canine<br>(This study) | OK523385.1<br>13598 bp<br>Feline<br>(This study) | NC_021145.1<br>AB732959.1<br>14296 bp<br>Canine<br>(Nakao et al., 2013) | MN099047.1<br>14226 bp<br>Canine<br>(Xie et al., 2019) | MG587892.1<br>13598 bp<br>Feline (Labuschagne et al., 2018) |
|---|---|---|---|---|---|
| **OK523384.1<br>14296 bp<br>Canine<br>(This study)** | 100.00% | 84.26% | 99.82% | 97.65% | 84.17% |
| **OK523385.1<br>13598 bp<br>Feline<br>(This study)** | 84.26% | 100.00% | 84.25% | 86.21% | 99.51% |


Labuschagne, M., Beugnet, F., Rehbein, S., Guillot, J., Fourie, J., Crafford, D., 2018. Analysis of Dipylidium caninum tapeworms from dogs and cats, or their respective fleas - Part 1. Molecular characterization of Dipylidium caninum: genetic analysis supporting two distinct species adapted to dogs and cats. Parasite 25, 30.

Nakao, M., Lavikainen, A., Iwaki, T., Haukisalmi, V., Konyaev, S., Oku, Y., Okamoto, M., Ito, A., 2013. Molecular phylogeny of the genus Taenia (Cestoda: Taeniidae): proposals for the resurrection of Hydatigera Lamarck, 1816 and the creation of a new genus Versteria. Int J Parasitol 43, 427-437.

Xie, Y., Liu, Y., Gu, X., Meng, X., Wang, L., Li, Y., Zhou, X., Zheng, Y., Zuo, Z., Yang, G., 2019. Complete mitogenome of the dog cucumber tapeworm. Mitochondrial DNA B Resour 4, 2670-2672.

Table S2. Cestode species, GenBank accession numbers and numbers of genes used in the BUSCO analysis. There were 128 ortholog genes present in all genomes listed

| **Species** | **GenBank Accession for genomes** | **Number of complete and single-copy ortholog genes** |
|---|---|---|
| ***Hymenolepis microstoma*** | GCA_000469805.3 | 619 |
| ***Hymenolepis nana*** | GCA_900617975.1 | 560 |
| ***Hymenolepis diminuta*** | GCA_900708905.1 | 602 |
| ***Spirometra erinaceieuropaei*** | GCA_902702965.1 | 501 |
| ***Echinococcus multilocularis*** | GCA_000469725.3 | 511 |
| ***Echinococcus granulosus*** | GCA_000524195.1 | 513 |
| ***Taenia asiatica*** | GCA_001693035.2 | 490 |
| ***Taenia saginata*** | GCA_001693075.2 | 501 |
| ***Taenia solium*** | GCA_001870725.1 | 484 |
| ***Taenia multiceps*** | GCA_001923025.3 | 509 |
| ***Schistocephalus solidus*** | GCA_017591395.1 | 539 |
| ***Moniezia expansa*** | GCA_019097775.1 | 531 |
| ***Echinococcus canadensis*** | GCA_900004735.1 | 515 |
| ***Mesocestoides corti*** | GCA_900604375.1 | 550 |
| ***Dipylidium caninum* (China)** | GCA_017562135.1 | 603 |

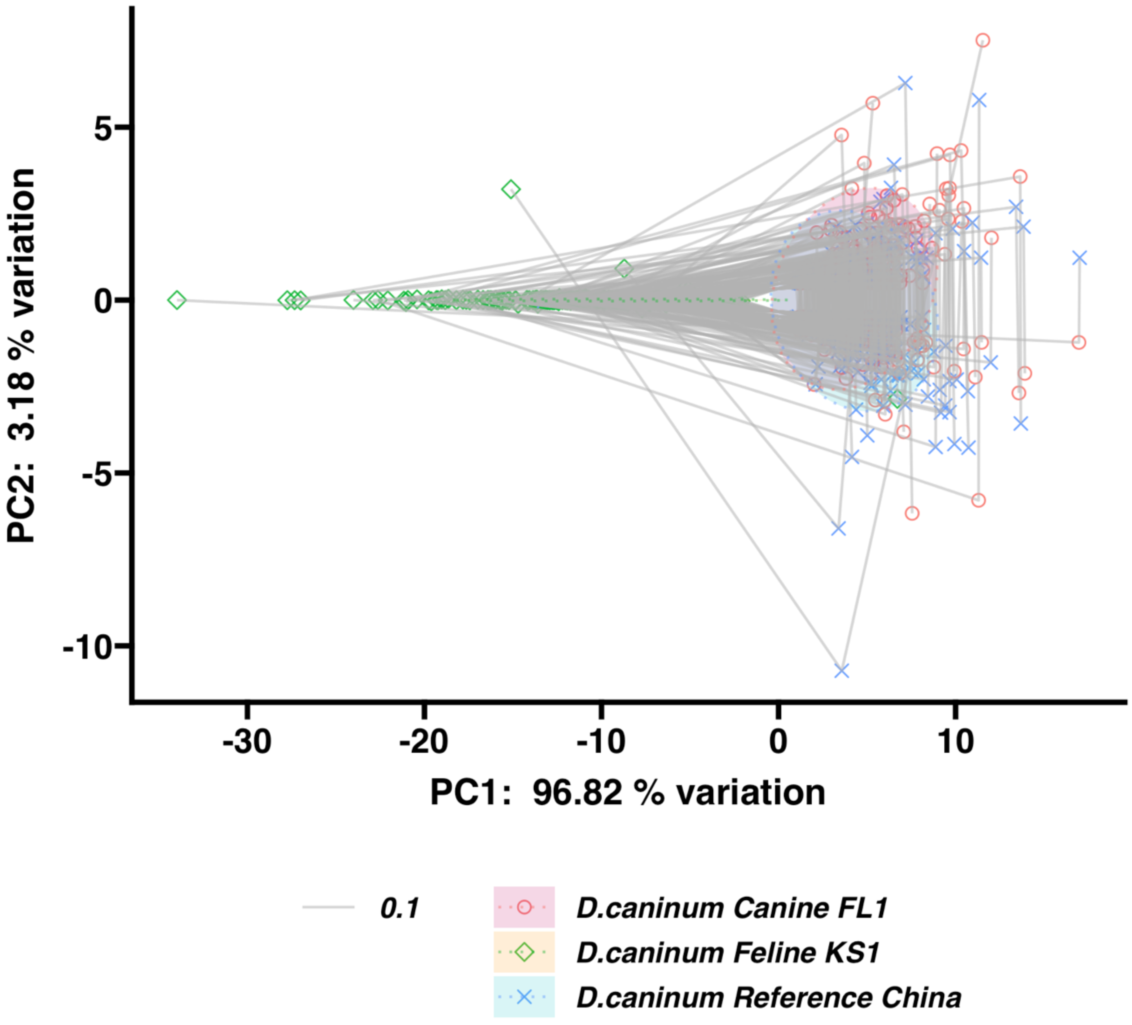


Figure S1. PCA showing the connected positions of components at each gene for each genome.

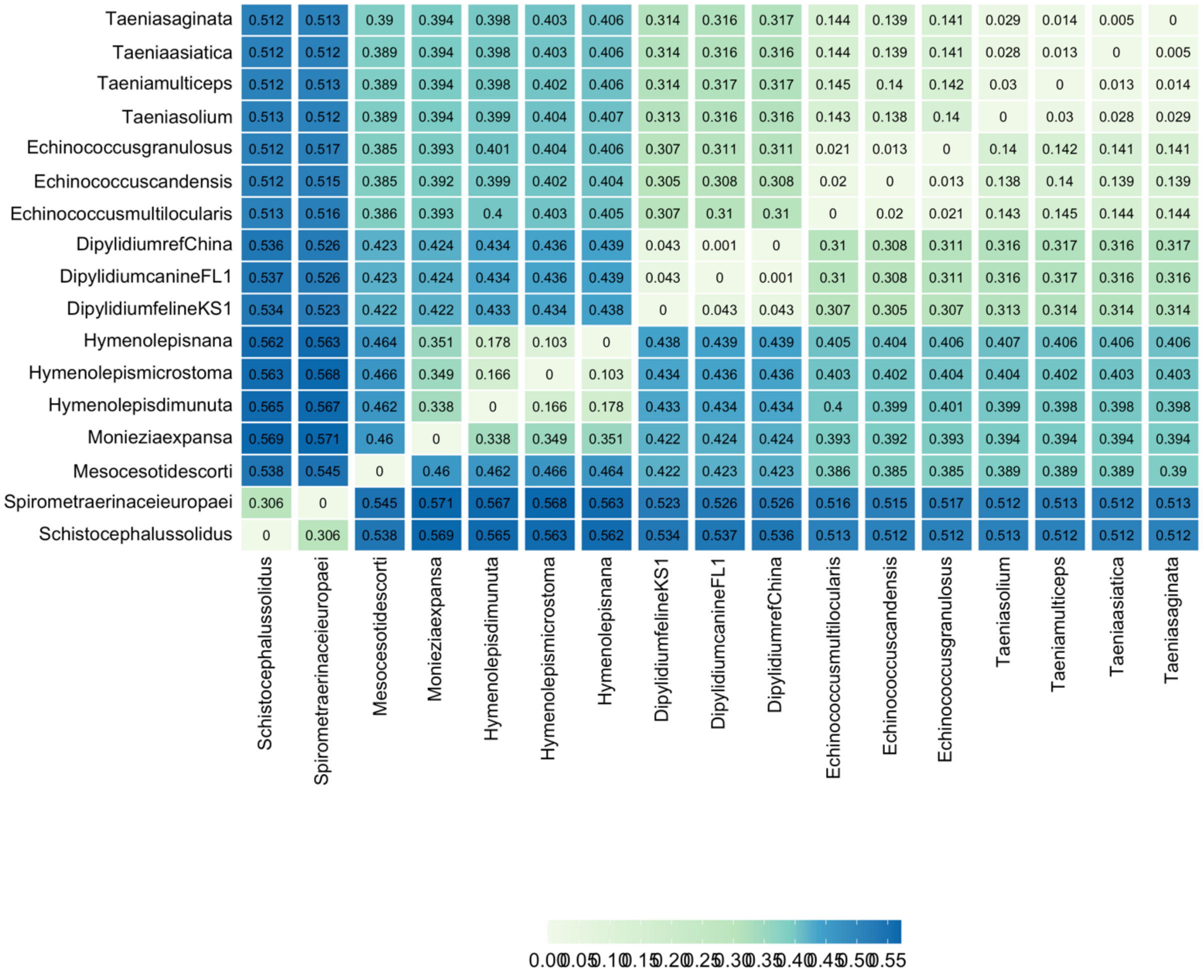


Figure S2. Pairwise genetic distances (Tamura-Nei, 1990) of the 128 gene supermatrix of BUSCOs from cestode species.

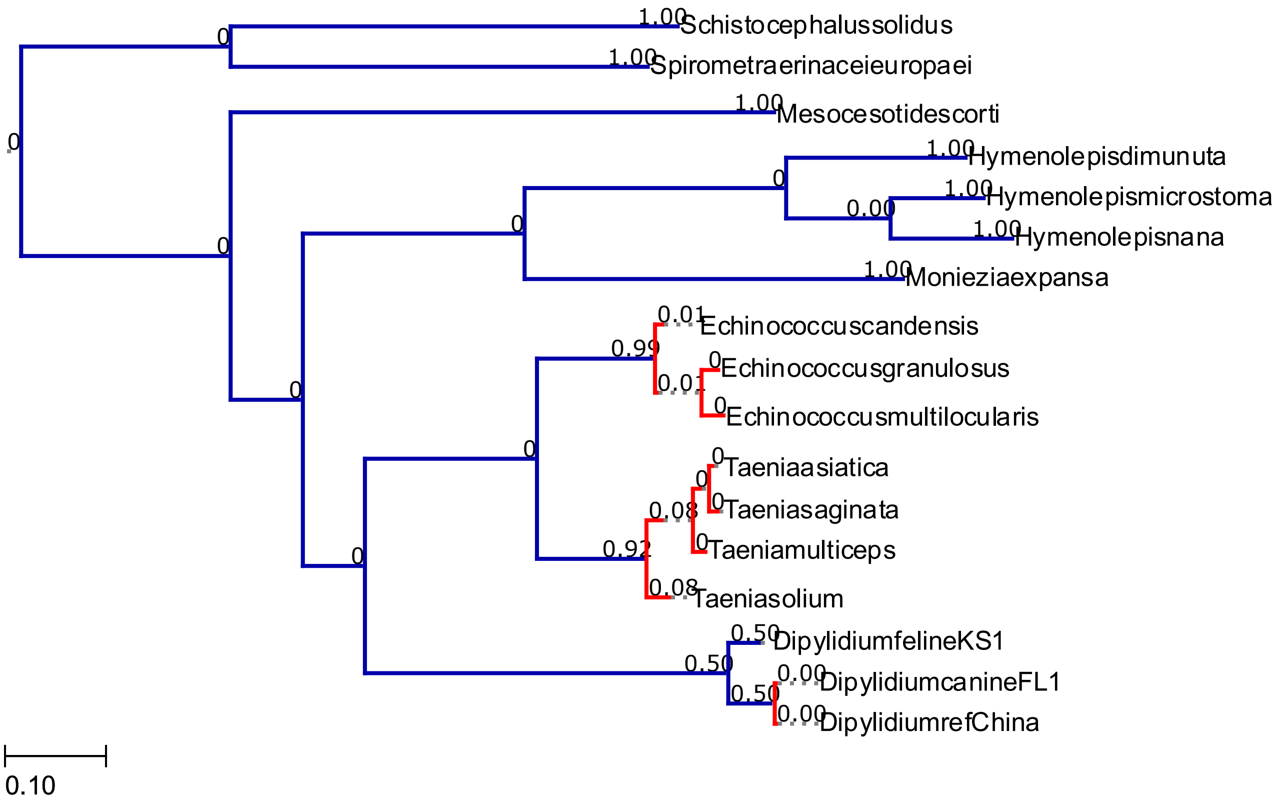


Figure S3. Output of bayesian PTP analysis of the ML phylogenetic tree created with a partitioned supermatrix 128 BUSCO genes in IQTree (See Figure 7). The tree was rooted on the Diphyllobothridean outgroup, with 100,000 MCMC generations, 100 thinning and 0.1 burn-in.

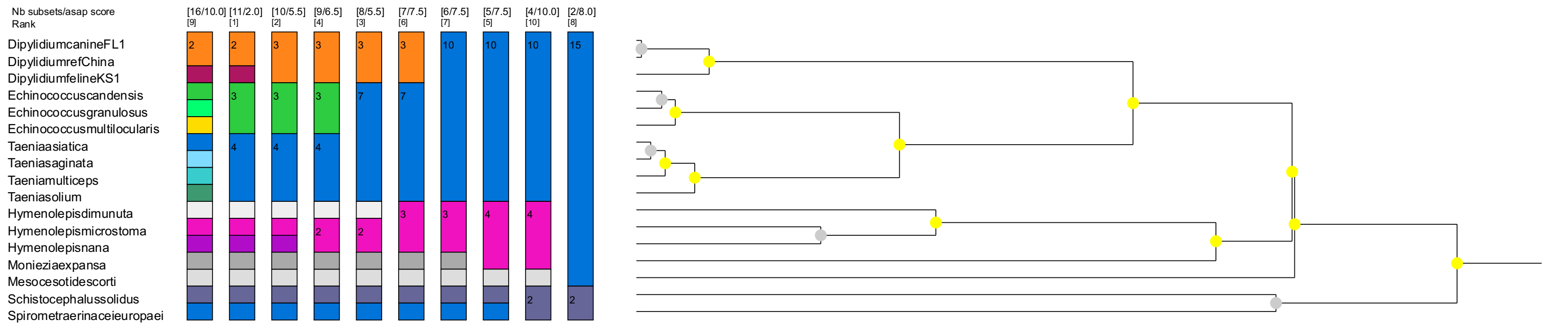


Figure S4: ASAP species delimitation output created with a fasta supermatrix of 128 BUSCO genes and Kimura 80 (Ti/Tv) substitution model.

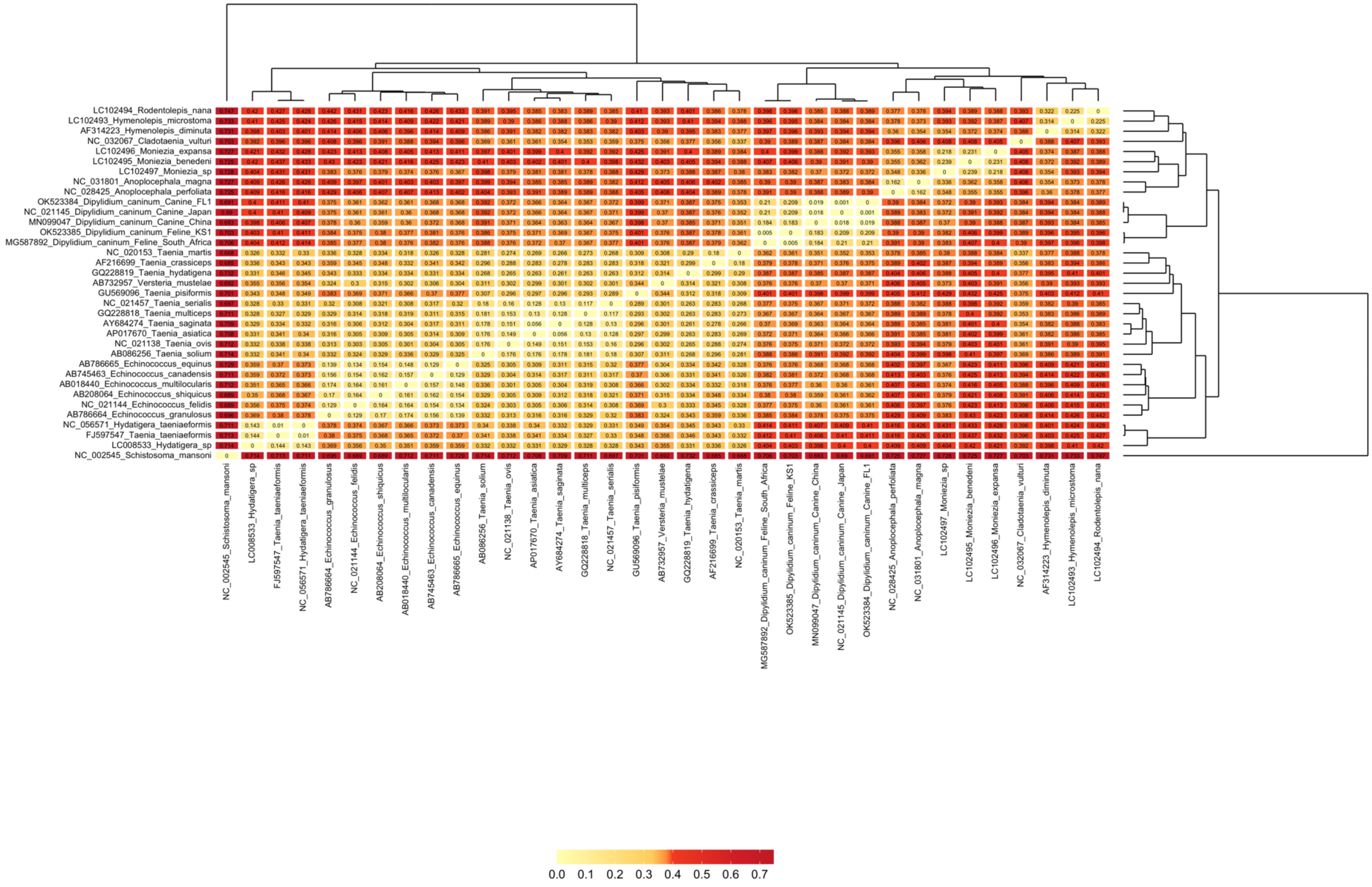


Figure S5. Pairwise genetic distances (Tamura-Nei, 1990) between concatenated 12 protein coding mitochondrial gene datasets.

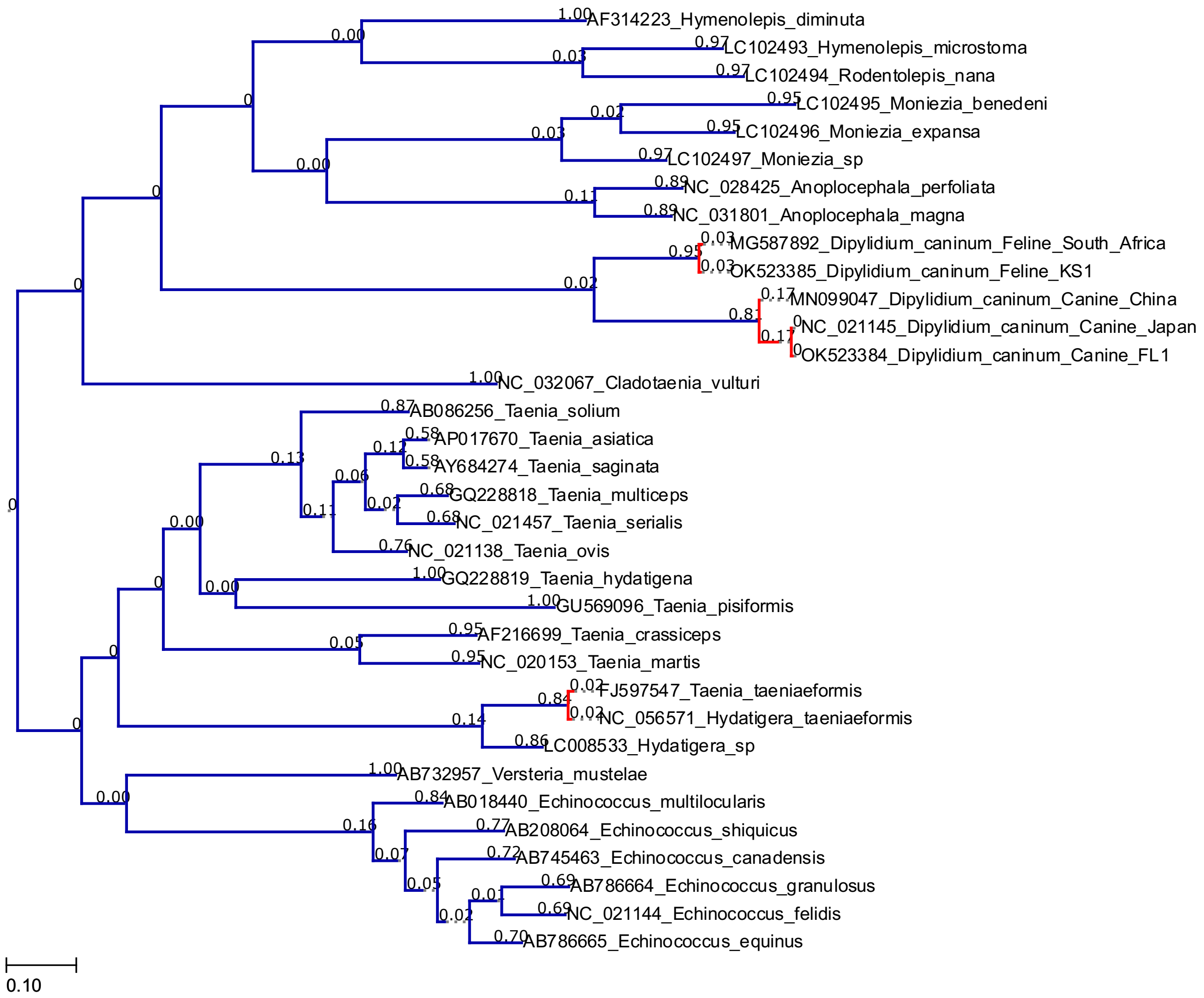


Figure S6. Output of bayesian PTP analysis of the ML phylogenetic tree created with partitioned mitochondrial 12 protein-coding nucleotide supermatrix in IQTree (See Figure 8). The tree was rooted on *Schistosoma mansoni* and the outgroup was removed to improve delimitation. Analysis was performed with 100,000 MCMC generations, 100 thinning and 0.1 burn-in.

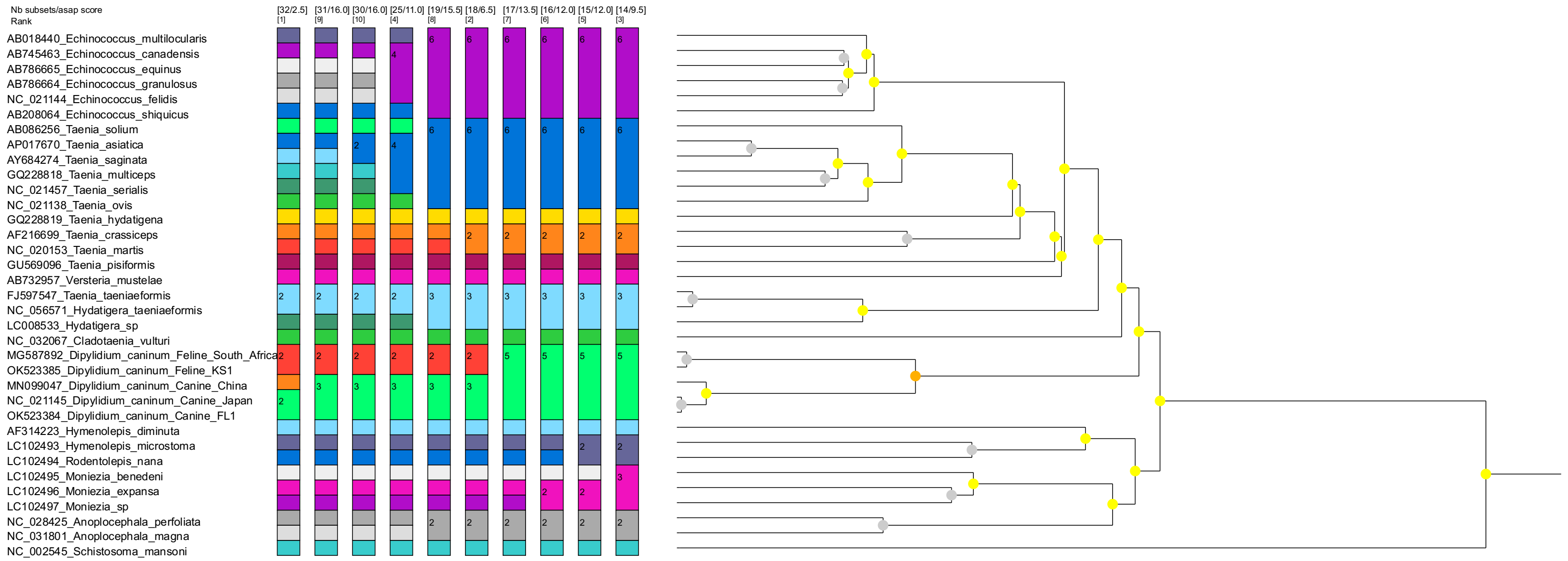


Figure S7: ASAP species delimitation output created with a fasta supermatrix of mitochondrial 12 protein-coding genes and Kimura 80 (Ti/Tv) substitution model.

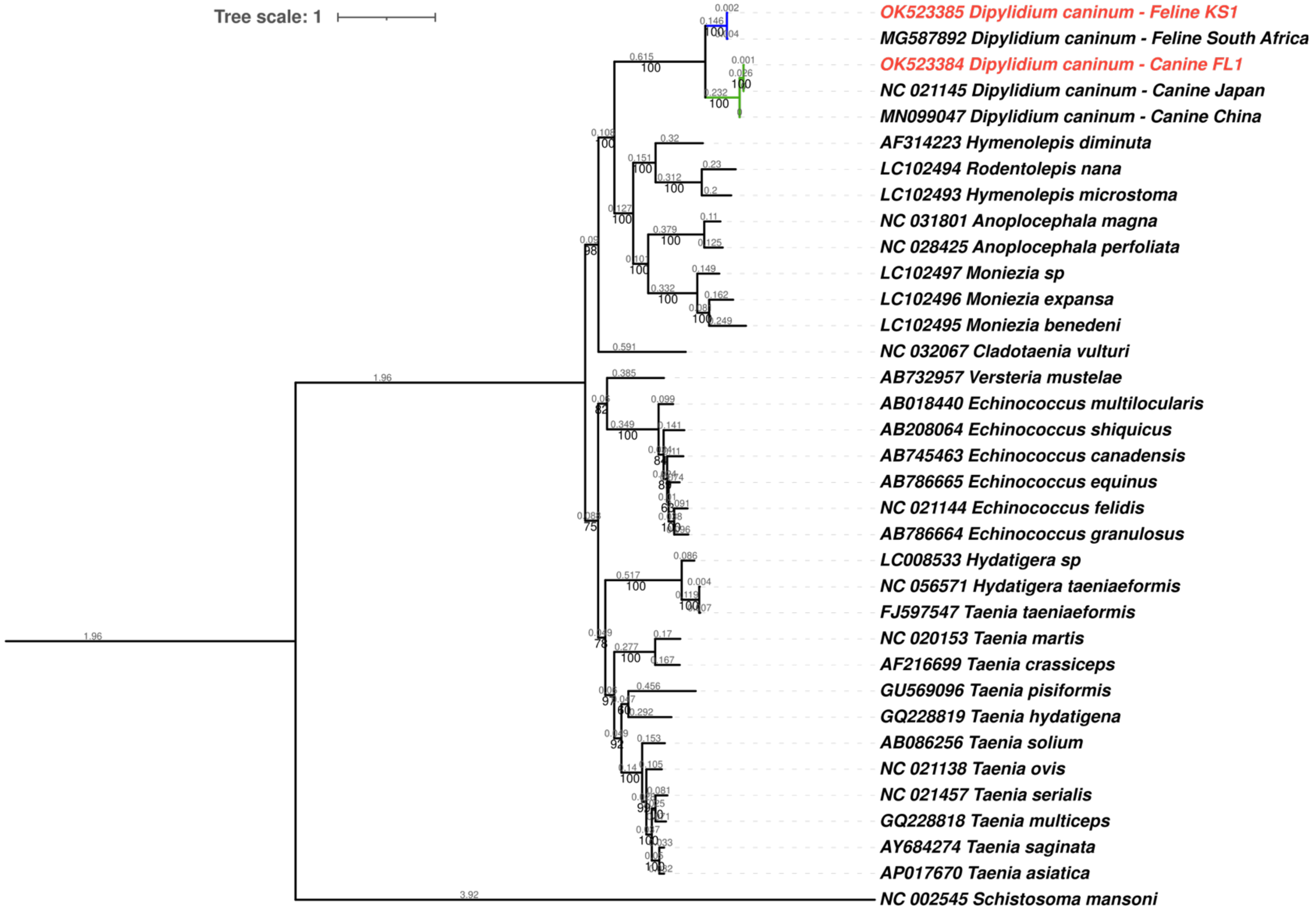


Figure S8. Maximum likelihood nucleotide phylogenetic tree of 12 mitochondrial protein coding genes of *Dipylidium caninum* mitochondrial genomes with branch lengths is available.
